# GRAYU: Graph-based Database integrating Ayurvedic formulations, medicinal plants, phytochemicals and diseases

**DOI:** 10.1101/2025.10.26.684703

**Authors:** Sarthak Joshi, Aditi Pathak, Dheemanth Reddy Regati, Revathy Menon, Deepthi S Ajith, Aditya Sheshadri, Neerja Viswanathan, Poulomi Ray, Vimarishi Koul, Pratishruti Panda, Shriya Anand Bhambore, Shailya Verma, Ananya Sinha, K. Mohamed Shafi, Murugavel Pavalam, Ramanathan Sowdhamini

**Author notes:** Joint corresponding authors (Murugavel Pavalam, Ramanathan Sowdhamini). These authors contributed equally to this work and share first authorship (Sarthak Joshi, Aditi Pathak, Dheemanth Reddy Regati).

## Abstract

The translation of India’s extensive traditional knowledge on indigenous medicinal plants into modern therapeutic solutions is contingent upon a systematic framework. While traditional Indian medicine offers a rich source of therapeutic leads, this knowledge is often not structured for modern computational analysis, creating a barrier to systematic drug discovery. To this end, we present GRAYU, a curated and comprehensive online database that integrates data across multiple categories, connecting 1039 traditional formulations to 12,949 indigenous plants, 129,542 phytochemicals, and 13,480 indicated diseases. It also provides insights into 1,382,362 plant-phytochemical, 116,824 plant-disease, 2,405 plant-formulation, and 4,087 formulation-disease associations. Case studies on phytochemical analogs, sustainable plant substitution, and disease-network pharmacology showcased the potential of integrative graphs to decode shared molecular signatures and therapeutic networks across traditional Ayurvedic formulations. GRAYU represents a user-friendly resource for researchers to investigate complex bio-associations and formulate novel therapeutic hypotheses, with insights from traditional Indian medicine. GRAYU is available at https://caps.ncbs.res.in/GRAYU/.

## 1 Introduction

Traditional medical systems such as Ayurveda, practiced in India for more than 3,000 years, constitute a vast repository of empirical knowledge on medicinal plants, polyherbal formulations, and their therapeutic applications (Gurib-Fakim 2006; *Charaka*). Despite this long history and widespread usage, much of the information remains dispersed across classical texts, ethnobotanical surveys, and modern phytochemical studies, limiting its accessibility and integration into contemporary biomedical research. As natural products continue to play a critical role in drug discovery, with approximately 34% of approved small-molecule drugs in the last four decades being natural products (Newman and Cragg 2020), natural product derivatives, or botanical drugs, there is a pressing need to digitize and standardize traditional medicine knowledge for computational and translational use.

In recent years, several databases have been developed to collate information on medicinal plants and their phytochemicals. For example, the IMPPAT 2.0 database provides a manually curated resource of over 4,000 Indian medicinal plants, nearly 18,000 phytochemicals, and more than 1,000 therapeutic uses. It offers chemical structures (2D and 3D), physicochemical and drug-likeness scores, and ADMET predictions (Vivek-Ananth et al. 2023). Similarly, OSADHI (Kiewhuo et al. 2023) is another platform aiming to digitize traditional Indian medicine, compiling phytochemical data for Indian medicinal plants and enabling chemoinformatics methods in natural-product drug development. Beyond the Indian traditional medicine sphere, integrative efforts like SymMap (Wu et al. 2019) have sought to combine Traditional Chinese Medicine (TCM) with modern medicine through symptom mapping and molecular mechanism links. Similarly, BATMAN-TCM (Liu et al. 2016; Kong et al. 2024) provides predicted TCM ingredient–target protein interactions, while HERB (Fang et al. 2021; Gao et al. 2025) integrates clinical trials and meta-analyses with experimental evidence for TCM. These efforts highlight the growing importance of databases that connect traditional formulations to modern molecular and clinical data.

Despite these advances, important gaps remain. Many existing platforms emphasize plant–compound or plant–therapeutic associations, but do not systematically incorporate classical polyherbal formulations, which are central to Ayurveda. In addition, Ayurvedic disease terms, often recorded in Sanskrit or regional languages, lack direct mapping to standardized biomedical vocabularies, creating barriers to integration with global biomedical resources. Finally, most existing interfaces provide limited functionality for multi-layered exploration, restricting users to linear searches rather than enabling graph-based traversals across plants, compounds, formulations, and diseases.

To address these limitations, we present GRAYU, a graph-based resource for Ayurveda that integrates over 12,000 medicinal plants, more than 1000 Ayurvedic formulations, nearly 130,000 phytochemicals, and over 13,000 diseases into a unified framework. GRAYU maps Ayurvedic nosology to modern disease ontologies to enable interoperability, while providing interactive graph visualization and advanced search pipelines to support complex queries. By combining curated associations with computational tools, GRAYU not only preserves traditional knowledge but also provides a platform for drug discovery and mechanistic exploration.

## 2 Materials and methods

### 2.1 Data curation

The GRAYU database was constructed by integrating extensive data from multiple sources, covering Ayurvedic formulations, medicinal plants, phytochemicals, and diseases. The curation process involved several key stages to ensure data accuracy, consistency, and comprehensive integration.

### 2.2 Curation of Formulations

Formulations were obtained from the Ayurvedic Standard Treatment Guidelines, the Ayurvedic Pharmacopoeia of India and the Ayurvedic Formulary of India. The Ayurvedic Standard Treatment Guidelines are available on the website of the National Ayush Mission, Ministry of Ayush, Government of India, while the Ayurvedic Pharmacopeia and Formulary are available on the website of the Pharmacopeia Commission for Indian Medicine & Homeopathy, Ministry of Ayush, Government of India. While the Ayurvedic Standard Treatment Guidelines is the primary source of the data that maps formulations to diseases, the Ayurvedic Pharmacopeia and the Ayurvedic Formulary are the predominant sources for mapping the formulations to their constituent plants. The integration between the sources was done by assigning a unique standardised name based on the transliteration of the Sanskrit name, and a single node was created for multiple instances of the same formulation across both sources. Data integration in cases such as multiple mentions of a formulation, across sources, as well as standardising the transliteration of the Sanskrit names of the formulations, was done manually.

### 2.3 Curation of Phytochemicals and Medicinal Plants

#### 2.3.1 Phytochemical Data Consolidation

The phytochemical data was aggregated from several specialized databases, including CMAUP (Zeng et al. 2019; Hou et al. 2024), FooDB (“FooDB,” n.d.), HMDB (Wishart et al. 2007, 2022), IMPPAT (Mohanraj et al. 2018; Vivek-Ananth et al. 2023), and NPASS (Zeng et al. 2018; Zhao et al. 2023). To effectively integrate the information from these varied sources, we established PubChem (Kim et al. 2025) as the foundational reference for consolidation.

First, a comprehensive mapping was created to link phytochemical records from the various source databases to their corresponding PubChem Compound IDs (CIDs). The core of our phytochemical dataset was then initialized using detailed records from PubChem. Subsequently, a script systematically processed each phytochemical entry from the source databases. For each entry, relevant data fields were extracted based on a predefined template that specified the path to the information within the original JSON structure.

To maintain data integrity, properties from the source databases were only added to a phytochemical’s record if that property was not already present from the PubChem data. This approach ensured that the high-quality, standardized information from PubChem was preserved while enriching it with additional data from other specialized resources. Values were cleaned to remove extraneous characters, and in cases where a property contained a list of values, they were concatenated into a single string. The final, consolidated phytochemical data was then written to a CSV file, forming the phytochemical nodes of our knowledge graph.

#### 2.3.2 Medicinal Plant Data Integration

Data for medicinal plants was curated and unified from three primary databases: CMAUP, IMPPAT, and OSADHI. The integration process was designed to create a single, comprehensive record for each unique plant species.

A script iterated through the JSON files from each of the plant databases. The scientific name of the plant was designated as the primary identifier for unifying records from different sources. For each plant, a master record was created, and properties from the various databases were progressively merged.

A predefined schema guided the extraction and merging process, distinguishing between unique properties (e.g., scientific name, taxonomic IDs), where only a single value is expected, and list-based properties (e.g., synonyms, common names), where multiple values are permissible. For list-based properties, values from different sources were appended and delimited by a semicolon. A specific function was also implemented to clean and standardize the scientific names to a consistent format (e.g., “Genus species”). This meticulous process of extraction, cleaning, and merging resulted in a unified dataset of medicinal plants, which was then exported to a CSV file to serve as the plant nodes in the graph. Furthermore, the script also parsed and extracted relationships to diseases and phytochemicals, which were written to separate edge files for constructing the knowledge graph.

### 2.4 Curation of Disease Nodes and Ontologies

Disease ontologies were obtained from two primary sources: the Medical Subject Headings (MeSH; (Dhammi and Kumar 2014)) database (https://www.nlm.nih.gov/mesh/) and the Disease Ontology (DOID; (Baron et al. 2024)) via the Ontology Lookup Service (McLaughlin et al. 2025) (https://www.ebi.ac.uk/ols4/ontologies/doid). From MeSH, only entries under the Diseases branch (MeSH Tree ID: C) were included, with each unique identifier treated as a distinct disease node. Similarly, disease and symptom identifiers from the DOID ontology were extracted and converted into corresponding nodes. To avoid redundancy, nodes were merged wherever DOID provided mappings to MeSH terms.

Two types of intra-category relationships were incorporated: “IS_SUBCLASS_OF” and “HAS_SYMPTOM.” Parent-child hierarchies were derived from the MeSH and DOID ontology trees, while “has symptom” relationships were taken directly from DOID. Finally, Ayurvedic descriptions of diseases and conditions were manually mapped to the most closely corresponding disease nodes, based on semantic similarity between Ayurvedic terminology and the standardized MeSH or DOID labels.

### 2.5 Knowledge Graph Construction

The GRAYU database is implemented using Neo4j, a native graph database, to effectively model and query the complex relationships between Ayurvedic formulations, medicinal plants, phytochemicals, and diseases. The construction process involved defining a graph schema with constraints and indexes, followed by the systematic loading of nodes and the creation of relationships from the curated CSV files. The core schema consists of four primary node labels: Phytochemical, Plant, Disease, and Formulation. To ensure data integrity and optimize query performance, unique constraints were established for primary identifiers: the PubChem CID for Phytochemical nodes, the standardized scientific name for Plant nodes, a unique disease node id for Disease nodes, and the formulation name for Formulation nodes. Additionally, indexes were created on frequently searched properties, such as compound names, MESH and DOID identifiers, to accelerate data retrieval. Data was loaded into the graph in batches using the apoc.periodic.iterate procedure for efficient, parallelized processing. Each row from the respective CSV files was used to create a node with its properties meticulously mapped and type-cast (e.g., to integer, float, or boolean). Text fields containing multiple values delimited by semicolons were converted into list properties within the nodes. Once the nodes were loaded, relationships were created to connect them, forming the knowledge graph:

1. FOUND_IN: A relationship from a Phytochemical node to a Plant node, established by matching the phytochemical’s CID and the plant’s scientific name.
2. ASSOCIATED_WITH_DISEASE: A relationship from a Plant node to a Disease node, created by matching plants to MESH or DOID identifiers listed in the curated edge files.
3. IS_INGREDIENT_IN: A relationship from a Plant node to a Formulation node, with properties such as parts and quantity attached to the relationship itself.
4. ASSOCIATED_WITH: A relationship from a Formulation node to a Disease node, also including the corresponding ayurvedic_term as a property on the relationship.
5. IS_SUBCLASS_OF and HAS_SYMPTOM: Intra-disease relationships created by linking Disease nodes to each other based on the hierarchical and symptomatic information curated from the MESH and DOID ontologies.

This structured approach results in a robust and queryable knowledge graph, providing a comprehensive digital framework for exploring the multifaceted data of traditional Indian medicine.

### 2.6 Online Database construction

The online database was constructed using a backend server built with the Flask web framework in Python. A data access layer was developed to encapsulate all database interactions, translating high-level application logic-such as paginated data retrieval for browsing, execution of simple and complex multi-step advanced searches, and fetching detailed information for individual entities into optimized Cypher queries executed via the official Neo4j Python driver. The front-end was constructed with standard web technologies, including HTML for structure, CSS for styling, and JavaScript for dynamic interactivity. For data visualization, graph-specific endpoints provide data formatted for the Cytoscape.js library, enabling users to interactively explore the network of relationships between entities.

## 3 Results

### 3.1 Statistics of the Graph

The comprehensive data integration process culminated in the creation of GRAYU, a large-scale knowledge graph comprising 157,010 nodes and 1,520,687 relationships (edges). The phytochemicals constitute the largest set of nodes, underscoring the chemical diversity captured within the database. The most prevalent relationship is FOUND_IN (Phytochemical-Plant), followed by ASSOCIATED_WITH_DISEASE edges (Plant-Disease), representing the therapeutic links between plants and diseases. Other relationships, such as ASSOCIATED_WITH (Formulation-Disease) and IS_INGREDIENT_IN (Plant-Formulation), establish the core therapeutic framework of Ayurveda, while hierarchical relationships like SUBCLASS_OF provide ontological depth to the disease classifications.

The overall architecture of these connections is visually summarized in the meta-graph shown in (Figure 1). The schematic illustrates the primary node types and the directed edges that define the relationships between them, such as a plant being an ‘ingredient in’ a formulation, which in turn ‘is associated with’ a disease. This high-level view provides a clear conceptual framework of the GRAYU database, representing how multilayered traditional knowledge has been structured and interconnected within our system.

**Figure 1.**
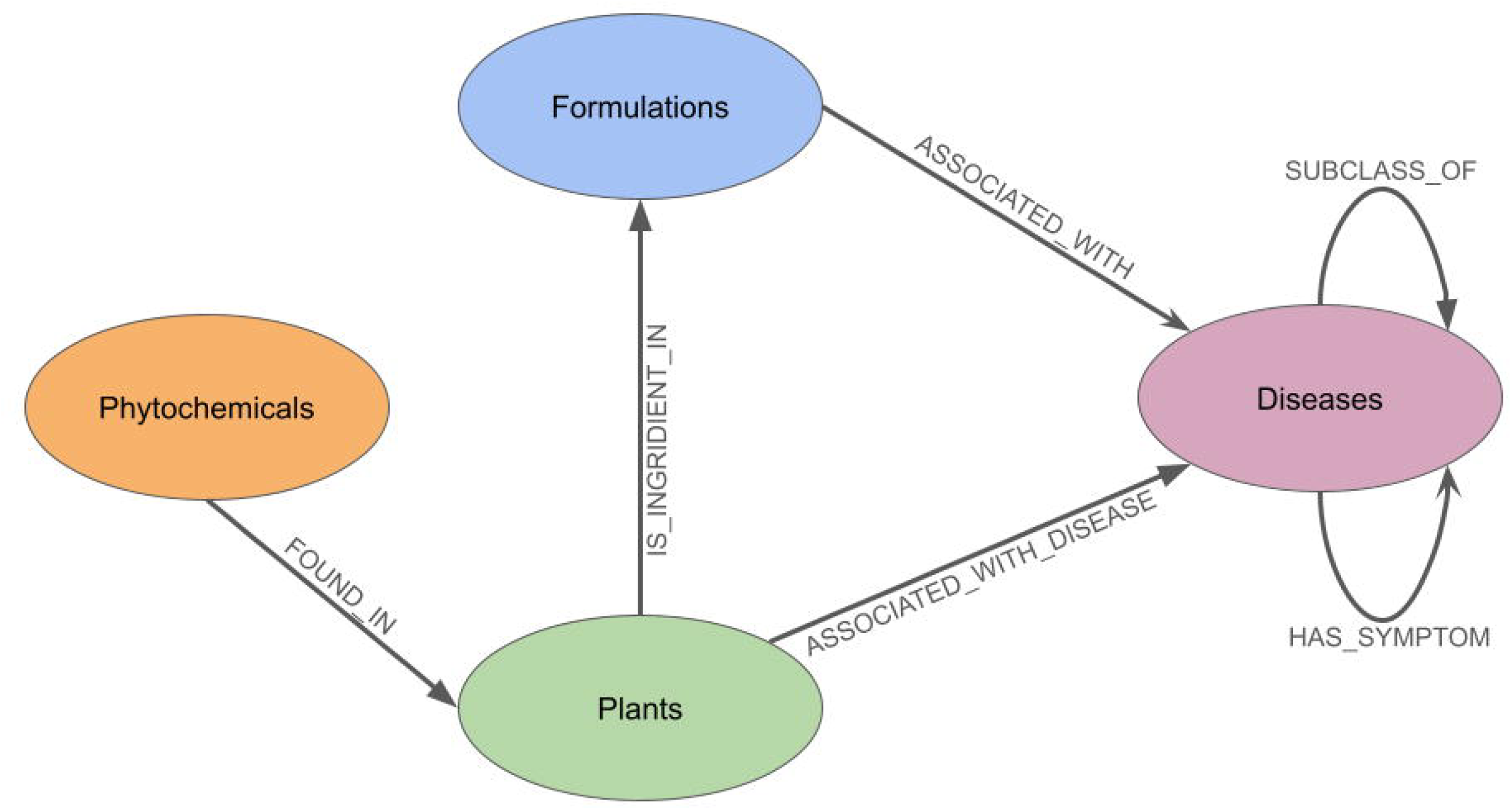
The meta-graph of the database which shows the types of nodes present, and the relationships which exist between them.

### 3.2 Website features and use cases

The GRAYU web portal provides a user-friendly interface for querying, browsing, and visualizing the integrated knowledge graph (Figure 2). The platform is designed to support a range of enquiries, from simple keyword searches to complex, multi-step queries, facilitating the exploration of relationships between formulations, plants, phytochemicals, and diseases.

**Figure 2.**
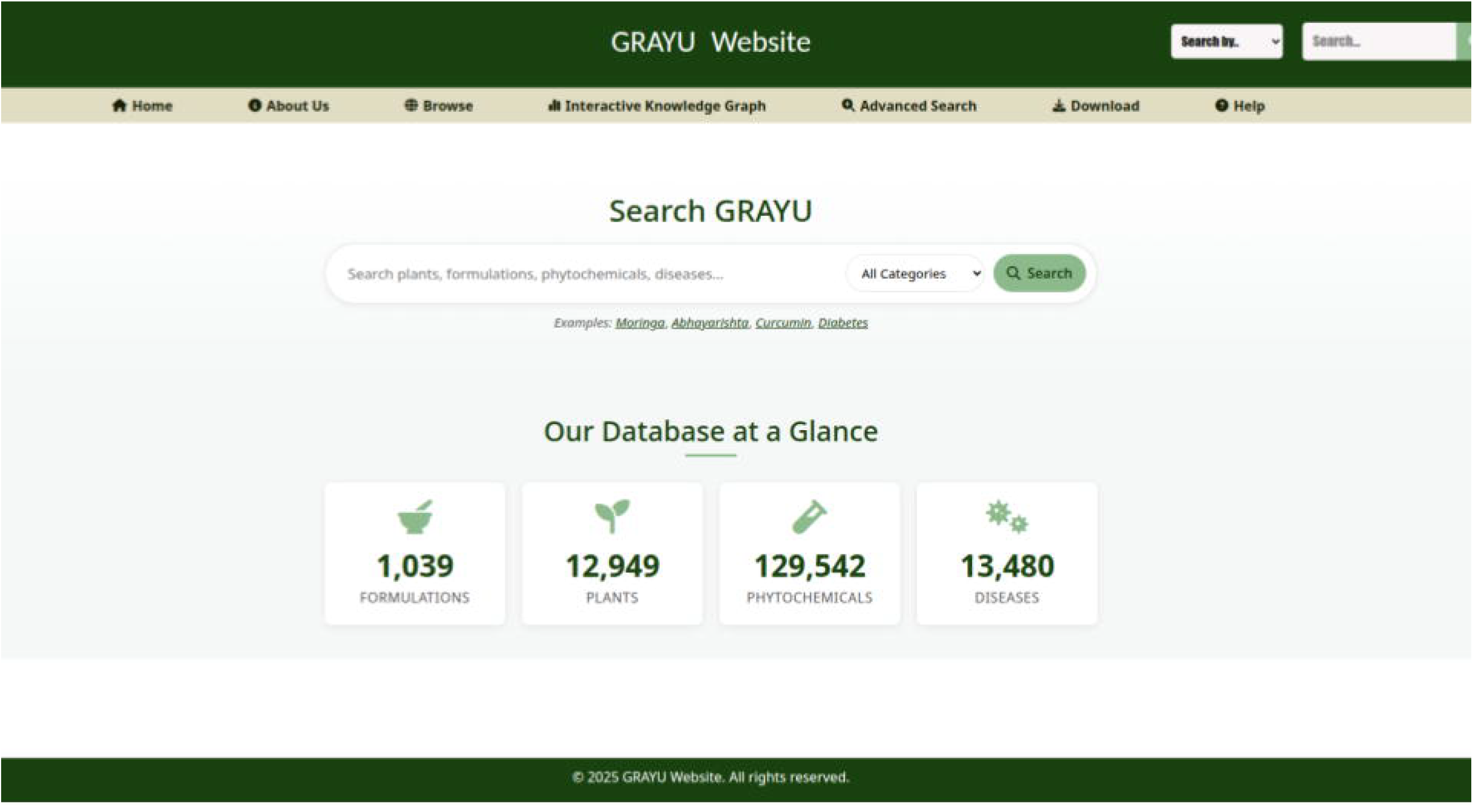
The homepage of the GRAYU web portal, presenting a navigation bar, a search bar and summary statistics of the database.

#### 3.2.1 Data Access and Search Functionalities

Users can access the data through several mechanisms. A quick search bar, available throughout the site, allows for keyword searches across all four categories (Formulations, Plants, Phytochemicals, and Diseases) and accepts scientific names, common names, and synonyms. Alternatively, a browse functionality presents the data in paginated, sortable tables for each category. Each entity in the database has a dedicated detailed results page, which presents its specific properties and associated connections. For example, a formulation page displays its ingredients and indicated diseases, while a plant page lists its constituent phytochemicals, associated diseases, and inclusion in formulations. These pages also feature an interactive graph visualization of the entity’s immediate network connections. For more complex inquiries, an advanced search page enables the construction of multi-step, filtered queries. Users can build a search pipeline where the results from one query serve as the input for the next, allowing for the progressive filtering and exploration of interconnected data. This feature supports the application of multiple filters based on the properties of both the primary entities and their connected nodes.

#### 3.2.2 Interactive Network Visualization

A dedicated graph page offers a powerful tool for dynamic network exploration. Users can search for any entity and render it as a central node in an interactive graph. The initial display can be customized to show a specific number of connections, which can then be dynamically expanded. Right-clicking on any node reveals options to expand its connections, either by a specific category (for instance, to show all phytochemicals for a plant) or by showing all of its relationships. A properties panel provides detailed information for any selected node, and the interface includes standard controls for navigation and visualization management.

### 3.3 Case Studies Demonstrating Applications of GRAYU

To illustrate the versatility of GRAYU, three case studies are presented, each highlighting a distinct application: (i) identification of molecular analogs through docking and cross-species mapping, (ii) selection of sustainable substitutes for threatened plants, and (iii) network pharmacology analysis of Ayurvedic formulations for Anemia. Together, they demonstrate the potential of GRAYU to integrate traditional pharmacology with molecular, ecological, and systems-level evidence.

#### 3.3.1 Molecular analogs of *Moringa oleifera* phytochemicals

*Moringa oleifera* is well recognized for its hypoglycemic and antioxidant properties attributed to bioactive constituents such as quercetin, kaempferol, and benzylamine (Shafi et al. 2022). To explore the structural analog landscape of these phytochemicals, molecular docking was performed against the diabetes-associated enzyme dipeptidyl peptidase-IV (DPP-IV), starting from the phytochemicals of Moringa and their analogs as captured from GRAYU. Among nearly 200 compounds screened, 4-hydroxybenzylamine, an analog of benzylamine, exhibited the strongest predicted interaction (-7.923 kcal/mol), surpassing the parent compound (Moringine or benzylamine with a docking score of - 6.937 kcal/mol), indicating enhanced affinity and potential inhibitory activity (Figure 3a; please see Supplementary Table 1 for full list). The *in silico* analysis showed the phytochemicals binding to the DPP-IV active site via distinct yet overlapping binding poses (Figure 4). GRAYU graph traversal revealed the co-occurrence of these metabolites in other plants such as *Sinapis alba* (white mustard) and *Brassica oleracea* (wild cabbage). Interestingly, GRAYU records both plants with anti-diabetic potential, reinforcing the validity of these molecular linkages (Gupta et al. 2022; Aboulthana et al. 2025). Despite these promising computational and graph-based findings, AI-assisted literature evaluation confirmed that no clinical or experimental evidence currently supports the therapeutic use of 4-hydroxybenzylamine for diabetes yet.. Nevertheless, this case highlights the GRAYU ability to connect molecular analogs across species and provides a data-driven route for rational exploration of novel or understudied compounds derived from Ayurvedic sources.

**Figure 3.**
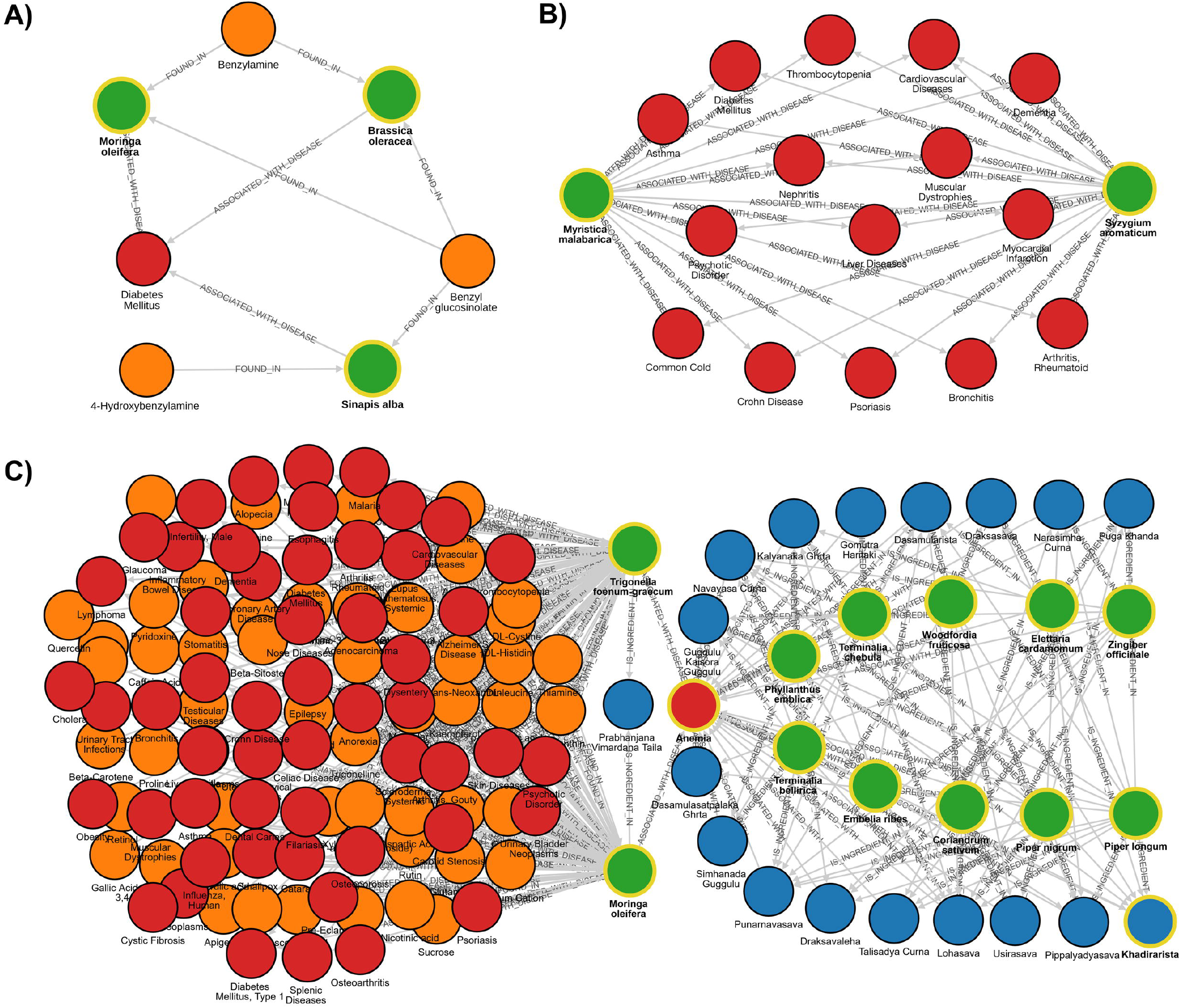
GRAYU graph-based case studies illustrating phytochemical, plant, and disease interconnections. (A) Molecular analogs of Moringa oleifera phytochemicals: network showing the relationship between benzylamine and its analog 4-hydroxybenzylamine, and their shared occurrence in Moringa oleifera, Sinapis alba, and Brassica oleracea, all linked to Diabetes Mellitus. (B) Replacement strategy for threatened species: graph depicting the biochemical and therapeutic overlap between Myristica malabarica (vulnerable species) and Syzygium aromaticum, sharing multiple phytochemicals and 15 common disease associations including arthritis, asthma, cardiovascular disorders, and diabetes. (C) Network pharmacology of anemia: multi-layered network connecting 17 Ayurvedic formulations (blue), 10 hub plants (green), associated diseases (red) and phytochemical (orange). The network highlights molecular convergence between Moringa oleifera and Trigonella foenum-graecum and identifies shared antioxidant and hematinic phytochemicals underlying anti-anemic formulations.

**Figure 4.**
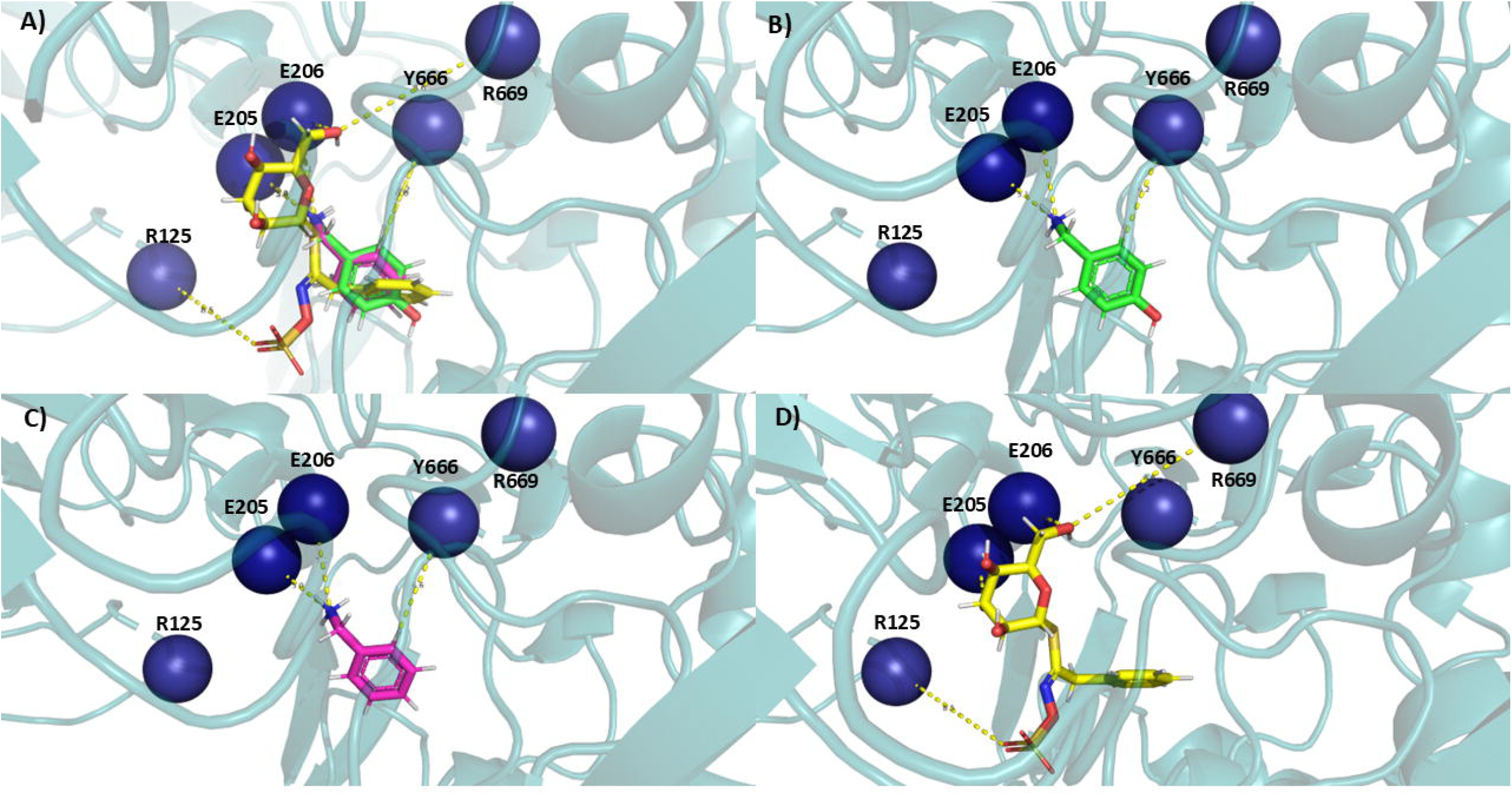
Molecular Docking Visualization of Phytochemicals within the DPP-IV Active Site. A) A comparative view of three phytochemicals of interest from Moringa oleifera are shown, their spatial overlap within the binding pocket as well as specific binding interactions with the protein are highlighted. The enzyme has been shown as a blue cartoon, the key residues are displayed as blue spheres and the interactions are shown as yellow dashed lines. B) Shows the binding pose of 4-hydroxybenzylamine (coloured green) highlighting interactions with the enzyme. C) Shows the binding pose of benzylamine (coloured pink) and D) shows the binding pose of Benzyl Glucosinolate (coloured yellow).

#### 3.3.2 Replacement strategy for threatened medicinal species

In Ayurveda, substitution of medicinal plants (*Abhava Pratinidhi Dravya*) is practiced when a herb becomes scarce or unavailable. Using GRAYU, we examined the substitution pair *Myristica fragrans* (nutmeg) and *Syzygium aromaticum* (clove) closely, where the aril of *M. fragrans* (mace) is traditionally replaced by *S. aromaticum* (Venkatasubramanian, Kumar, and Nair 2010; Menthe and Menthe 2017). GRAYU searches identified 155 shared phytochemicals, including eugenol, isoeugenol, myristicin, elemicin, 4-terpineol, camphene, β-caryophyllene, quercetin, kaempferol, ursolic acid, stigmasterol, dehydrodieugenol, p-cymene, β-phellandrene and linalool and 58 common disease associations, encompassing metabolic, inflammatory, and infectious disorders. While *M. fragrans* is listed as data deficient on the IUCN Red List, its wild relative *Myristica malabarica* is classified as vulnerable, highlighting the need for sustainable alternatives. Comparative graph exploration revealed 66 shared phytochemicals and 15 overlapping disease associations between *M. malabarica* and *S. aromaticum*, validating the latter as a promising substitute based on both biochemical and therapeutic similarity (Supplementary Table 2; Figure 3b). This case underscores GRAYU potential for conservation-driven substitution analysis, enabling evidence-based selection of ecologically viable alternatives without compromising on therapeutic equivalence. However, detailed comparisons of each feature, such as dosage levels, need to be carefully considered for replacement purposes.

#### 3.3.3 Network pharmacology of Anemia

Two medicinal plants, also used as traditional foods, *Moringa oleifera* and *Trigonella foenum-graecum* are popularly used for diabetes management (Pasha et al. 2020; Naika et al. 2022). Next, we demonstrate how GRAYU reveals biochemical and therapeutic convergence across these two taxonomically distinct Ayurvedic plants. Both plants are widely utilized in traditional medicine for metabolic and hematinic disorders and are rich in overlapping flavonoids and phenolic acids such as quercetin, kaempferol, rutin, β-sitosterol, and ferulic acid. These metabolites are well documented for their antioxidant, anti-inflammatory, and glucose-lowering activities, supporting the traditional co-usage of these plants in formulations targeting diabetes, arthritis, obesity, and anemia. The integrative graph of GRAYU highlighted that these shared phytochemicals create a common pharmacological axis linking *Moringa* and *Trigonella* through mutual disease associations, suggesting that chemically distinct species can yield functionally redundant or synergistic therapeutic effects (Figure 3c). This overlap supports the Ayurvedic concept of therapeutic equivalence and indicates potential synergistic or complementary use of the two herbs in multi-target formulations addressing metabolic and inflammatory disorders.

Extending from this plant-pair intersection to a disease-centric network, Anemia (Pandu Roga) was selected as a representative condition to explore multi-herbal connectivity. GRAYU identified 17 Ayurvedic formulations associated with anemia, comprising nearly 90 unique medicinal plants traditionally prescribed for *Rakta-vardhana* (blood enrichment). Network centrality analysis revealed ten recurrent hub species, including *Piper longum, Terminalia chebula, Phyllanthus emblica, Terminalia bellirica*, and *Zingiber officinale*, that form the structural backbone of anti-anemic formulations (Supplementary Table 3). These plants possess well-established hematopoietic, antioxidant, and Rasayana properties, experimentally validated for enhancing iron absorption, erythropoiesis, and oxidative stress regulation. Further expansion to phytochemical nodes revealed 20 core metabolites, notably β-sitosterol, α-tocopherol, gallic acid, palmitic acid, stearic acid, and rutin, that collectively define a shared biochemical scaffold mediating hematinic and metabolic homeostasis. The network demonstrates that Ayurvedic formulations for Anemia are systematically organized molecular ecosystems, designed to act through synergistic modulation of metabolic and redox pathways. GRAYU, thus, provides a computational framework to visualize, quantify, and rationalize these cross-layer interactions, bridging traditional therapeutic logic with systems pharmacology.

## 4 Discussion

The GRAYU database provides a unified framework for systematically connecting the diverse layers and categories of Ayurvedic knowledge with contemporary biomedical ontologies. The integration of over 12,000 medicinal plants, nearly 130,000 phytochemicals, more than 1000 Ayurvedic formulations and over 13,000 diseases in one platform represents one of the largest resources to date focused on traditional Indian medicine. The scale and breadth of the associations, spanning plant– phytochemical, plant–formulation, formulation–disease and plant–disease connections, highlight the potential of Ayurveda as a knowledge system that can be interrogated using modern computational tools.

In comparison with other existing resources, GRAYU expands the landscape of natural product databases in important ways. Earlier efforts such as IMPPAT 2.0 (Vivek-Ananth et al. 2023) and OSADHI (Kiewhuo et al. 2023) provided comprehensive collections of Indian medicinal plants and phytochemicals, but their focus remained largely at the level of plant–compound associations without embedding them within the context of classical formulations or mapping to standardized disease vocabularies. Similarly, global resources such as HMDB (Wishart et al. 2022) and NPASS (Zhao et al. 2023) have provided metabolite- and activity-centric perspectives, while Chinese medicine repositories including SymMap (Wu et al. 2019), BATMAN-TCM 2.0 (Kong et al. 2024), HERB 2.0 (Gao et al. 2025), and ETCM v2.0 (Zhang et al. 2023) have created powerful examples of integrating traditional systems with modern targets and diseases. GRAYU complements and extends these approaches by creating a graph-based architecture that is specific to Ayurveda, and explicitly bridges Sanskrit nosology with MeSH and DOID disease ontologies. This mapping is particularly important as it enables interoperability between Ayurvedic descriptions and widely adopted biomedical standards, thereby opening avenues for cross-database analyses and translational research.

The statistical analysis of GRAYU demonstrates the richness of the network, revealing a highly interconnected system where many plants contain phytochemicals shared across multiple formulations, and where a single disease node may be targeted through numerous plant–formulation combinations. This property reflects the holistic and polyherbal principles of Ayurveda, while at the same time, creating opportunities for network pharmacology approaches to identify convergent mechanisms.

A major strength of GRAYU lies in its user interface and graph-based exploration features. The implementation of Interactive Knowledge graph and advanced search pipelines allows researchers to construct complex, multi-step queries, tracing the relationships from plants to phytochemicals and onwards to diseases or formulations. The expandable graph visualizations provide an intuitive view of the immediate neighbourhood of a node, while the advanced filtering enables detailed interrogation of properties. Such functionality transforms GRAYU from a static catalogue into a dynamic exploratory platform with intuitive dialogues. This is aligned with the evolving expectations of modern bioinformatics resources, where the ability to interrogate complex associations is valued as highly as data comprehensiveness.

The GRAYU-based case studies collectively emphasize the capability to interlink traditional pharmacological wisdom with molecular and ecological data. The *Moringa oleifera* analog study demonstrates the ability to pinpoint structural derivatives and cross-species analogs that may possess improved pharmacodynamic potential, serving as entry points for rational natural product modification leading to rational drug design. The substitution analysis between *Myristica* and *Syzygium* highlights the utility of GRAYU in identifying molecularly and therapeutically equivalent alternatives, supporting conservation and sustainable sourcing. Traditional foods and superfoods may belong to taxonomically diverse lineages, yet converge in chemical resources and in enabling better disease management. This is shown by comparing *Moringa* and *Trigonella*. Finally, the anemia network model shows that Ayurvedic polyherbal systems are not arbitrary aggregates, but systematically optimized biochemical ensembles, with recurring phytochemicals providing a unifying molecular framework. Collectively, these case studies reveal that GRAYU enables systems-level exploration of Ayurveda, supporting applications from drug discovery and sustainability planning to mechanistic hypothesis generation. By integrating graph-based visualization with experimental and clinical annotations, GRAYU bridges traditional and modern pharmacology, offering a reproducible computational platform for future integrative medicine research.

There are, however, limitations to be acknowledged. The extent of coverage of knowledge captured within GRAYU in the four categories critically depends on the information arising from primary data resources. The graph networks necessarily follow a path across the categories - for example, from Plant-to-Disease and not from Phytochemical-to-Disease directly (as shown in Figure 1). The diversity of preparation methods and variation in formulation proportions cannot be fully captured in the current dataset. The mapping of Ayurvedic disease categories to modern ontologies, while carefully curated, remains an approximation in cases where concepts are not directly comparable. Moreover, the majority of associations remain at the level of traditional knowledge and computational inference, underscoring the need for systematic experimental validation. Despite these challenges, GRAYU represents a significant advance by creating a scaffold upon which such validations can be prioritised and guided.

In conclusion, GRAYU establishes a large-scale, curated, and graph-enabled platform that integrates data across categories (Ayurvedic formulations, plants, phytochemicals, and diseases) into a unified framework. By combining data richness, ontology mapping, advanced search functionality, and illustrative case studies, GRAYU not only preserves traditional knowledge in digital form but also positions it for active use in modern biomedical research and drug discovery.

## 6 Author Contributions

MP and RS conceptualized the study. MP, SJ, AP, DR, SAB, DSA and AS integrated data. RM, NV, PR, VK, PP, SV, AS and MS performed data analysis and visualization. SJ, AP, DR, SAB, DSA, AS, RM, NV, PR, VK, PP, SV AS and MS wrote the manuscript. MP and RS wrote the final draft of manuscript.

## 8 Acknowledgments

Authors would like to acknowledge Prof. Sudhir Krishna, Drs Adwait Joshi, Abhishek Sharma and Meenakshi Iyer for discussions. We thank the vibrant and helpful discussions from various Ayurvedic practitioners and formulators, especially Dr. Rajesh Sreenivasan of BV Pundit Sadvaidyasala, Nanjangud and Drs. Subrahmanya and Vishnuprasad of TransDisciplinary University, Bangalore, which helped to curate the database. We thank NCBS (TIFR) for infrastructural facilities.

## Supplementary Material

Supplementary Information is available in the supplementary file.

## Data Availability Statement

This paper presents an analysis of existing, publicly available data. The GRAYU database is publicly available at https://caps.ncbs.res.in/GRAYU/. All source data underlying the findings are accessible through the database.

