## Supplementary File for "GRAYU: Graph-based Database integrating Ayurvedic formulations, medicinal plants, phytochemicals and diseases"

*Supplementary Table 1: Docking Results of Phytochemicals against DPP-IV enzyme.*

| **Phytochemicals present in *Moringa* /Analogs** | Molecular Docking Scores (kcal/mol) |
| --- | --- |
| **4-Hydroxybenzylamine** | **-7.923** |
| **Benzyl glucosinolate** | **-7.177** |
| (2R,3S,4S)-3-[(2S,3R,4R,5S,6R)-3,4-dihydroxy-6-(hydroxymethyl)-5-[(2S,3R,4S,5R,6R)-3,4,5-trihydroxy-6-(hydroxymethyl)oxan-2-yl]oxyoxan-2-yl]oxy-2-(3,4,5-trihydroxyphenyl)-3,4-dihydro-2H-chromene-4,5,7-triol | -7.145 |
| Glucoconringiin | -7.109 |
| Benzyl beta-d-glucopyranoside | -7.056 |
| **Benzyl glucosinolate** | **-6.998** |
| 4-(3'-O-acetyl-alpha-l-rhamnosyloxy) benzyl isothiocyanate | -6.971 |
| Benzylamine | -6.937 |
| Naringenin 7-O-(2'',6''-di-O-alpha-rhamnopyranosyl)-beta-glucopyranoside | -6.916 |
| Benzyl glucosinolate | -6.894 |
| Benzyl glucosinolate | -6.835 |
| Glucoconringiin | -6.786 |
| 2-[3-hydroxy-4-[(2R,3R,4R,5R,6S)-3,4,5-trihydroxy-6-methyloxan-2-yl]oxyphenyl]acetonitrile | -6.779 |
| Benzyl glucosinolate | -6.774 |
| Moringin | -6.766 |
| Isobutyl glucosinolate | -6.703 |
| Benzyl glucosinolate | -6.689 |
| Benzyl glucosinolate | -6.682 |
| Glucoconringiin | -6.666 |

*Supplementary Table 2: Common Disease Connections between Myristica malabarica and Syzygium aromaticum*

Psoriasis

Dementia

Psychotic Disorder

Common Cold

Bronchitis

Thrombocytopenia

Liver Diseases

Myocardial Infarction

Arthritis Rheumatoid

Nephritis

Asthma

Crohn Disease

Diabetes Mellitus

Muscular Dystrophies

Cardiovascular Diseases

*Supplementary Table 3: Hub plants (10) and associated phytochemicals (20) identified from Ayurvedic formulations (17) for anemia in GRAYU.*

| **No.** | **Formulations** | ***Piper longum*** | ***Terminalia chebula*** | ***Piper nigrum*** | ***Zingiber officinale*** | ***Phyllanthus emblica*** | ***Terminalia bellirica*** | ***Coriandrum sativum*** | ***Woodfordia fruticosa*** | ***Elettaria cardamomum*** | ***Embelia ribes*** |
| --- | --- | --- | --- | --- | --- | --- | --- | --- | --- | --- | --- |
| 1 | Dasamularista | Yes | Yes | – | – | Yes | Yes | Yes | Yes | Yes | Yes |
| 2 | Dasamulasatpalaka Ghrta | Yes | – | – | Yes | – | – | – | – | – | – |
| 3 | Draksasava | Yes | – | – | – | – | – | – | Yes | Yes | – |
| 4 | Draksavaleha | Yes | – | – | Yes | Yes | – | Yes | – | – | – |
| 5 | Gomutra Haritaki | – | Yes | – | – | – | – | – | – | – | – |
| 6 | Guggulu Kaisora Guggulu | Yes | Yes | Yes | Yes | Yes | Yes | – | – | – | Yes |
| 7 | Kalyanaka Ghrta | – | Yes | – | – | Yes | Yes | – | – | Yes | Yes |
| 8 | Khadirarista | Yes | Yes | – | – | Yes | Yes | – | Yes | Yes | – |
| 9 | Lohasava | Yes | Yes | Yes | Yes | Yes | Yes | Yes | Yes | – | Yes |
| 10 | Narasimha Curna | Yes | – | Yes | Yes | – | – | Yes | – | – | – |
| 11 | Navayasa Curna | Yes | Yes | Yes | Yes | Yes | Yes | – | – | – | Yes |
| 12 | Pippalyadyasava | Yes | – | Yes | – | – | – | – | Yes | Yes | Yes |
| 13 | Puga Khanda | Yes | – | Yes | Yes | – | – | Yes | – | Yes | – |
| 14 | Punarnavasava | Yes | Yes | Yes | Yes | – | Yes | Yes | Yes | – | – |
| 15 | Simhanada Guggulu | – | Yes | – | – | Yes | Yes | – | – | – | – |
| 16 | Talisadya Curna | Yes | – | Yes | Yes | – | – | – | – | Yes | – |
| 17 | Usirasava | – | – | Yes | – | – | – | Yes | Yes | – | – |
|  | **Phytochemicals** |  |  |  |  |  |  |  |  |  |  |
| 1 | Palmitic Acid | Yes | Yes | Yes | Yes | Yes | Yes | Yes | Yes | Yes | Yes |
| 2 | Beta-Sitosterol | Yes | Yes | Yes | Yes | Yes | Yes | Yes | Yes | Yes | Yes |
| 3 | Stearic Acid | Yes | Yes | Yes | Yes | Yes | Yes | Yes | Yes | Yes | Yes |
| 4 | Nicotinic Acid | Yes | Yes | Yes | Yes | Yes | – | Yes | Yes | Yes | Yes |
| 5 | Alpha-Tocopherol (Vit E) | Yes | Yes | Yes | Yes | – | Yes | Yes | Yes | Yes | Yes |
| 6 | Rutin | Yes | Yes | Yes | Yes | Yes | Yes | Yes | Yes | – | – |
| 7 | Linoleic Acid | – | Yes | Yes | Yes | Yes | Yes | Yes | Yes | Yes | – |
| 8 | Ascorbic Acid (Vit C) | – | Yes | Yes | Yes | Yes | Yes | Yes | Yes | Yes | – |
| 9 | Oleic Acid | – | Yes | Yes | Yes | Yes | Yes | Yes | Yes | Yes | – |
| 10 | Elaidic Acid | Yes | Yes | Yes | Yes | – | Yes | Yes | – | Yes | Yes |
| 11 | Caryophyllene | Yes | Yes | Yes | Yes | Yes | – | Yes | – | Yes | Yes |
| 12 | Isoquercetin | Yes | – | Yes | Yes | – | – | Yes | Yes | – | Yes |
| 13 | Eucalyptol | Yes | – | Yes | Yes | – | – | Yes | Yes | Yes | – |
| 14 | Triacontane | Yes | Yes | – | Yes | – | Yes | Yes | Yes | – | – |
| 15 | Kaempferol | Yes | – | Yes | Yes | Yes | – | – | Yes | – | Yes |
| 16 | Riboflavin (Vit B₂) | – | – | Yes | Yes | Yes | – | Yes | Yes | Yes | – |
| 17 | D-Galactose | Yes | – | Yes | Yes | – | Yes | Yes | Yes | – | – |
| 18 | Chinese Gallotannin | – | Yes | Yes | Yes | Yes | Yes | – | Yes | – | – |
| 19 | Gallic Acid | – | Yes | – | Yes | Yes | Yes | Yes | Yes | – | – |
| 20 | Myrcene | Yes | – | Yes | Yes | Yes | – | Yes | – | Yes | – |
